# Behavioural hyperactivity and risk-taking in young-adult male mice correlate with increased dopamine release in dorsal, but not ventral, striatum

**DOI:** 10.64898/2026.09.23.753631

**Authors:** Sean Coady, Anusha Kamesh, Christina Gentle, Bruno Di Marco Vieira, Camila Tiefensee-Ribeiro, Jeanne Madranges, Austen Milnerwood

## Abstract

Adolescence is characterized by increased novelty seeking, impulsivity, locomotion and risk-taking, behaviours that facilitate the transition to adulthood but may also increase vulnerability to psychiatric illness. Maturation of dopamine (DA) transmission within the striatum is thought to contribute to these behavioural changes. However, the developmental trajectory of striatal DA release remains unclear, with conflicting evidence regarding age- and sex-dependent changes, particularly across dorsal and ventral striatal regions. Complicating interpretation, many studies classify mice younger than 2 months as “adults,” potentially obscuring important developmental differences.

Here, we examined how age, sex and striatal subregion influence DA release during adolescence and early adulthood. DA release was measured in dorsal and ventral striatal brain slices from male and female C57BL/6J mice at 2 and 4 months of age, corresponding to adolescence and young adulthood. Ultrafast imaging of the genetically encoded DA sensor, dLight, was used to quantify extracellular DA transients evoked by low- and high-frequency electrical stimulation. Glutamatergic and cholinergic transmission was pharmacologically blocked to isolate DA release intrinsic to dopaminergic axons. DA release measures were related to exploratory and locomotor behaviour.

We identified a male-specific increase in dorsal striatal DA release between 2 and 4 months of age that coincided with increased locomotion and risk-taking. In contrast, DA release in the ventral striatum did not show comparable age- or sex-dependent changes. These differences were not attributable to altered glutamatergic or cholinergic contributions to DA release. Together, these findings demonstrate that maturation of striatal DA transmission is region- and sex-dependent, with prominent developmental changes occurring in the dorsal striatum during the transition from adolescence to adulthood. These findings help reconcile conflicting reports of developmental DA release and emphasize the importance of considering age, sex and striatal subregion when defining mature dopaminergic function in rodents.

**Research Highlights:**

- Ultrafast dLight imaging captures developmental changes in dopamine release
- Young adult males exhibit transiently elevated dorsal striatal dopamine release
- Elevated dopamine release correlates with hyperactivity and risk-taking
- Sex differences are intrinsic to release from dopamine axons
- Ventral striatal dopamine release remains predominantly stable across age and sex

**Graphical Abstract:** 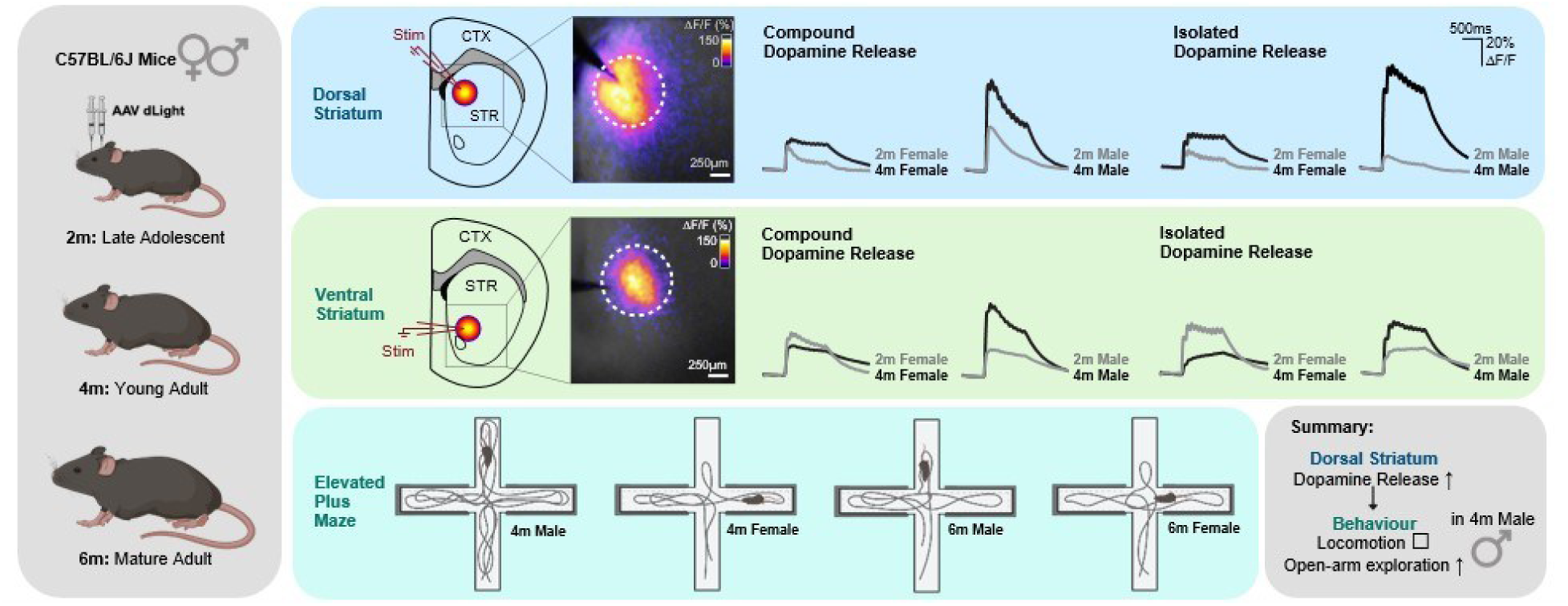

## Introduction

In humans and rodents, adolescence is marked by increased risk-taking, novelty seeking, hyperactivity and impulsivity (Adinolfi et al., 2018; Knoll et al., 2000). This is thought to support the transition to independent adulthood and facilitate the acquisition of new skills and adaptive behaviours, but also contributes to the elevated risk of psychiatric illness in adolescents (Adinolfi et al., 2018; Knoll et al., 2000; Paus et al., 2008; Spear, 2000, 2011, 2013; Steinberg, 2008). These developmental changes are thought to be driven in part by maturation of the dopaminergic system in the frontal cortex and in the basal ganglia, a group of subcortical nuclei involved in action selection, motor learning, skill acquisition, goal-directed and habitual behavior (Leisman et al., 2014; Redgrave et al., 2010; Yin & Knowlton, 2006).

The striatum is the largest nucleus and primary input structure of the basal ganglia, where glutamatergic sensory-motor, associative, and limbic input from many thalamic and all cortical areas are integrated by spiny projection neurons (SPNs). Striatal output through SPNs is heavily modulated by dopamine (DA) (Calabresi et al., 2014); DA is required for SPN maturation and also regulates glutamate synaptic release and plasticity (Calabresi et al., 2014; Lieberman et al., 2018; Sulzer et al., 2016). Additionally, di-synaptic axonal DA release is driven by glutamate activity via cholinergic interneurons, independent of ascending DA neuron action potentials (Kosillo et al., 2016; Threlfell & Cragg, 2011; Tritsch et al., 2012). The striatum is also functionally divided by the source of DA input. The dorsal striatum (DS), innervated by dopaminergic axons projecting from the substantia nigra pars compacta (SNc), is involved in goal-directed behaviour, habit learning, and movement selection (Bromberg-Martin et al., 2010; Faure et al., 2005; Redgrave et al., 2010; Yin & Knowlton, 2006). In contrast, DA innervation of the ventral striatum (VS) comes from the ventral tegmental area (VTA) and regulates motivation, novelty seeking, and reinforcement learning (Kim & Kaang, 2022; Palmiter, 2008; Salamone & Correa, 2012; Schultz et al., 1997; Threlfell & Cragg, 2011).

In mice, striatal DA axons reach morphological maturity by 2 weeks of age, and DA transmission regulates the maturation of SPN excitability over the first month of postnatal development (Lieberman et al., 2018). How DA release properties mature from juvenile, through adolescence, to adult ages is a subject of intense interest, but the literature is confounded by contradictory reports. In the DS, DA release was shown to increase between 1-4 months with no sex differences (Lieberman et al., 2018). However, other reports show that DS DA release is higher in males than females at 1 month (Brundage et al., 2022), or that it is comparable in both sexes at 1 month, and that female release reduces by 2 months (Pitts et al., 2020). In the VS, contradictory findings of both increased and decreased DA release have been reported across age and sex (Badanich et al., 2006; Brundage et al., 2022; Gonzalez et al., 2024; Iacino et al., 2024; Pitts et al., 2020; Stamford et al., 1989).

At 1-month of age, (C57BL/6J) mice begin puberty and display their first interest in sex, thus entering ‘adolescence’. ‘Onset of adulthood’ (i.e. adolescence) is at 2 months, ‘full adulthood’ is at 6, and ‘middle-aged adult” is over 8-15 months (Arellano et al., 2024). Several reports show rodents are hyperactive and exhibit enhanced novelty seeking and impulsivity during adolescence (Adinolfi et al., 2018; Knoll et al., 2000). A current problem in neuroscience is highlighted by a meta-analysis of 85 “adult” mouse neurogenesis studies, which found 49% used mice aged <2 months (adolescent) and only 7% included mice >4 months (Snyder, 2019). The premature assignment of adult status conflates early and late developmental stages with mature stages, if not entirely missing the latter.

To directly address the influence of age, sex, and DA transmission in striatal subregions, here DA release was assayed in brain slices prepared from male and female mice at 2 and 4 months of age, corresponding to adolescents and young adults (Arellano et al., 2024). Ultrafast imaging of the genetically encoded dopamine sensor, dLight, was used to quantify extracellular DA transients evoked by low- and high-frequency electrical stimulation. We isolated release intrinsic to DA axons by pharmacological blockade of glutamatergic and cholinergic transmission. Results were correlated to exploratory behaviour to determine if developmental changes to DA release correspond with activity levels and risk-taking.

We found a male-specific increase in DS DA release in young adults at 4 months, coincident with increased locomotion and risk-taking. Further, we identified that age- and sex-dependent differences in DA release were apparent in the dorsal, but not ventral striatum. These changes were not explained by differences in the contribution of glutamatergic & cholinergic inputs to DA release. The data help clarify existing literature and reveal sex-specific differences in striatal DA release throughout adolescence, into early adulthood.

## Methods

### Animals

Late adolescent (2m; P48-60), young adult (4m; P97-113), and mature adult (6m; P170-190) C57B16/J mice were housed and bred in accordance with the Canadian Council on Animal Care regulations (Animal Use Protocol 2017-7888B). All procedures were approved by and governed in accordance with the Neuro Centre of Neurological Disease Models (Animal Use Protocol 2017-7888B).

### Elevated Plus Maze

Behavioral testing was performed between 7:00 pm and 4:00 am (mouse awake cycle) by a single experimenter. All mice were transferred to the testing room 30min prior to testing to habituate. The EPM apparatus (Maze Engineers Inc. (USA) was in the center of a large metallic enclosure (2 x 2 x 2m) surrounded by dark curtains to avoid direct light. Video recordings were captured on a computer using the Motive 2.0 software (Optitrack, USA). Individual mice were placed in the center area of the maze after recording was started and moved freely for 10 minutes before recording was terminated. Following recording, mice were returned to their home cage. Between trials, the apparatus was cleaned with distilled water, 70% ethanol, then distilled water again. The center of the mouse body, as seen from above, was used as the point of reference for tracking by Ethovision XT 13 software (Noldus, USA). The number of entries and the time spent in each zone (open arm, closed arm, center), distance traveled, velocity, and transitions between arm types were recorded.

### Surgery

The modified dopamine D1 receptor fluorescent reporter (dLight1.3b) was expressed in the striatum by stereotaxic injection of adeno-associated virus (AAV) constructs (AAV5-CAG-dLight1.3b; Addgene, 125560-AAV5, 1.5×10^13 GC/mL). Mice were injected subcutaneously with carprofen (2-4 mg/mL, 20mg/kg) and allowed to rest for >15 mins. Isoflurane (with oxygen) was used to anesthetize the mice (5% for induction, 2% for maintenance) then placed in a stereotaxic head frame (Kopf Instruments) with eye lubrication provided before subcutaneous injection of Marcaine above the site of incision >3 mins prior to surgery. An incision was made in the scalp, and the skull levelled with bregma / lambda as reference. A craniotomy over the site of injection in each hemisphere (distance from bregma in mm: 1.0 lateral, 1.0 anterior, 4.1 ventral for VS injection, and 1.0 or 2.0 lateral, 1.0 anterior, -3.2 ventral for DS injection). A 10µL Hamilton-style syringe (Nanofil) attached to a microinjector (Harvard Apparatus Pump11 Elite) was lowered to coordinates after zeroing to the brain surface, then 300 nL of AAV was injected into the VS at 1nL/sec. After 5 minutes rest, the needle was moved up to DS, and a second injection of 300nL conducted at 1nL/sec followed by another 5-minute rest period. The needle was then moved to the opposite hemisphere, and injections were repeated. The needle was then withdrawn and the scalp rehydrated with Marcaine and closed with sutures (4-0 silk; Ethicon 683 G) and reinforced with Vetbond (3M 1469SB). Subcutaneous 0.9% NaCl injection was given to replace fluids (0.2–0.5mL/10 mg) and mice were monitored for pain and discomfort after regaining consciousness prior to return to home cage in ∼1h. Post-operative monitoring over 3 days included daily subcutaneous carprofen injection.

### Brain Slicing & dLight Imaging

Optical recordings of DA release were conducted in 300µm acute coronal slices, as previously described (Kamesh et al., 2025). Mice were restrained, decapitated, brains extracted and immediately placed in ice-cold recovery solution (1 min; in mM: 93 NMDG, 93 HCl, 2.5 KCl, 1.2 NaH_2_PO_4_, 30 NaHCO_3_, 20 HEPES, 25 glucose, 5 sodium ascorbate, 3 sodium pyruvate, 10 MgSO_4_, 0.5 CaCl_2_, pH 7.3-7.4, 290-310mOsm; carbogen-infused) for axonal DA isolation experiments, or artificial cerebrospinal fluid (aCSF; in mM: 125 NaCl, 3 KCl, 25 NaHCO_3_, 1.25 NaH_2_PO_4_, 2 MgCl_2_, 2 CaCl_2_, 10 glucose, pH 7.2-7.4, 290–310mOsm, carbogen-infused) for the remoxipride experiment. Striata were isolated by rostral and caudal blocking, then hemisected after attachment to the vibratome plate with cyanoacrylate (Leica Micro-systems VT 1200S). Coronal slices were cut at 0.22 mm/sec, immediately transferred to 35°C recovery solution or aCSF for 15 minutes, then transferred to a holding chamber perfused with oxygenated aCSF at 22-25°C for >45 min before recording in the same solution.

dLight recordings were obtained with a 4X objective, wide-field illuminated by blue light (473nm, CoolLED pE-340fura), under the control of a high-speed shutter, ultrafast EM-CCD camera (Andor iXon Ultra 897). Transient release was evoked by a stimulus isolator (WPI A365) triggered by Clampex (Digidata 1550B) connected to a monopolar tungsten stimulating electrode (A-M Systems 574000) lowered 50–100µM beneath the surface of the tissue. Recordings were captured with Andor Solis software (4 × 4 binning) at 205Hz. Spontaneous dLight transients were excluded from analysis of responses to electrical stimulation. Throughout, no-stimulation and stimulation trials were interleaved (1-minute interval) for background fluorescence bleach subtraction *post-hoc*. For compound vs isolated axonal DA release experiments, aCSF containing picrotoxin (PTX; 100µM) was perfused prior to isolation of intrinsic DA axon release with the addition of mecamylamine (MEC; 20µM), D-2-amino-5-phosphonopentanoate (D-AP5; 10µM), and 6-cyano-7-nitroquinoxaline-2,3-dione (CNQX; 10µM) for 15 minutes prior to resumption of recording. For D2 autoreceptor block, the D2R antagonist remoxipride hydrochloride (Remox, 10µM; VWR International Co. 77992-667), was perfused for a minimum of 15 minutes prior to pulse train experiments.

### Stimulation Paradigms

Low-frequency paired-pulse (0.2ms, 4s IPI) were delivered at increasing stimulation intensity with a 2-minute inter-sweep interval between each stimulus to generate input-output curves and gauge peak release. The stimulation intensity which evoked 50-60% maximal peak was chosen for pulse-train experiments. High-frequency pulse-trains were 10 pulses at 10Hz, with an 11th ‘recovery’ pulse administered at an increasing inter-pulse interval (500, 1000, 2500, and 5000ms) after the final pulse of the 10-pulse train. 4 repetitions of the pulse-trains were delivered with 2 minutes between each repetition. For isolation and D2 autoreceptor block experiments, pharmacology was added to the aCSF and perfused for >15 minutes prior to repeating the pulse-train protocol. To test D2 auto-receptor block, pulse-train protocols were followed by paired-pulse experiments matching the inter-pulse intervals of the recovery pulses.

### Data Presentation & Statistical Analyses

Data curation and statistics were conducted in GraphPad Prism11. Parametric data were analyzed using 1- or 2-way ANOVA, with Holm-Šídák multiple comparisons tests used for *post-hoc* analyses when ANOVA main effects reached significance (p < 0.05). Pairwise comparisons were performed using Fisher’s LSD test where indicated. Asterisks represent significance of pair-wise comparisons \**p* < 0.05, \*\**p* < 0.005, \*\*\**p* < 0.0005, \*\*\*\**p* <0.0001. Trends were defined as *p=*0.05 to 0.10. Statistical analyses are described in figure legends. Data are presented as n=observations from (n) animals (e.g., WT = 6(3) is 6 observations from 3 WT animals).

## Results

### Young adult male mice exhibit transiently increased exploratory and risk-tolerant behaviour

We compared male and female mice at 4 (young adult) and 6 (full adult) months of age in the elevated plus maze (EPM; Fig1). We found 4m males were hyperactive compared to age-matched females and 6m males, exhibiting higher overall distance travelled (Fig.1A) and more entries into the maze’s open arm (Fig.1B). As the number of entries into the closed arm were comparable between groups (Fig.1C). We interpret this as increased risk-taking behavior in young adult males, relative to full adults. The differences did not correlate with levels of blood estradiol in females or testosterone in males (Fig.S1). The data indicate that exploratory behaviour is elevated in young adult, but not full adult, male mice.

**Figure 1.**
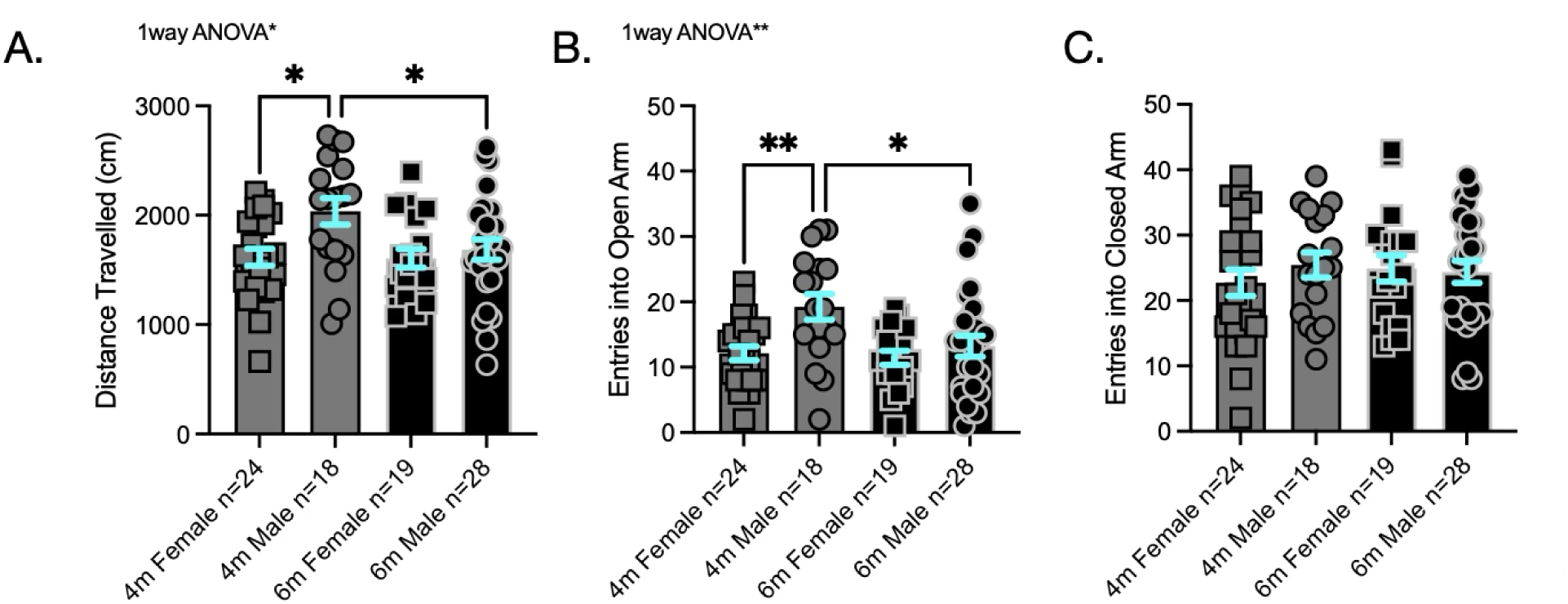
Exploratory behaviour is transiently elevated in young adult male mice. Exploratory behaviour in the elevated plus maze. **A-C)** 4m male mice travelled greater total distances (**A**; 1-way ANOVA F_(3, 85)_=3.99, *p*=0.01) and spent more time in the open arm (**B**; 1-way ANOVA F_(3, 85)_=4.88, *p*=0.004) than 4m females and 6m males. There was no difference in time spent in the closed arms between groups (**C;** 1-way ANOVA F_(3, 85)_=0.37, *p*=0.8). Holm-Šídák multiple comparisons test \**p*<0.05, \*\**p*<0.005.

### Dorsal striatal dopamine release is markedly elevated in 4-month-old males

To determine if behavioural differences in young adult male mice correlate with altered DA function, we quantified evoked release in acute brain slices from 4m male and female mice. We also compared adolescent males and females at 2m. Experiments were conducted in the presence of picrotoxin (PTX) to eliminate inhibitory gamma-aminobutyric acid (GABA)_A_ signalling. At both 2 and 4m, dLight transient peak fluorescence in the DS evoked by low frequency 2-pulse stimulation (4s IPI) was higher in males than females across the range of stimulation intensities (Fig.2A-D). Maximum dLight responses were higher in 4m males than 4m females and 2m males, indicating elevated male DA release at this age (Fig.2D). Conversely, paired-pulse ratios (PPRs; 4s IPI) were highest in 4m females (Fig.2E), suggesting lower initial release probability (Pr) and/or reduced D2 autoreceptor (D2AR) activity. Notably, there were no differences in DA transient decay rates, indicating no differences in DA re-uptake between groups (Fig.S2).

**Figure 2.**
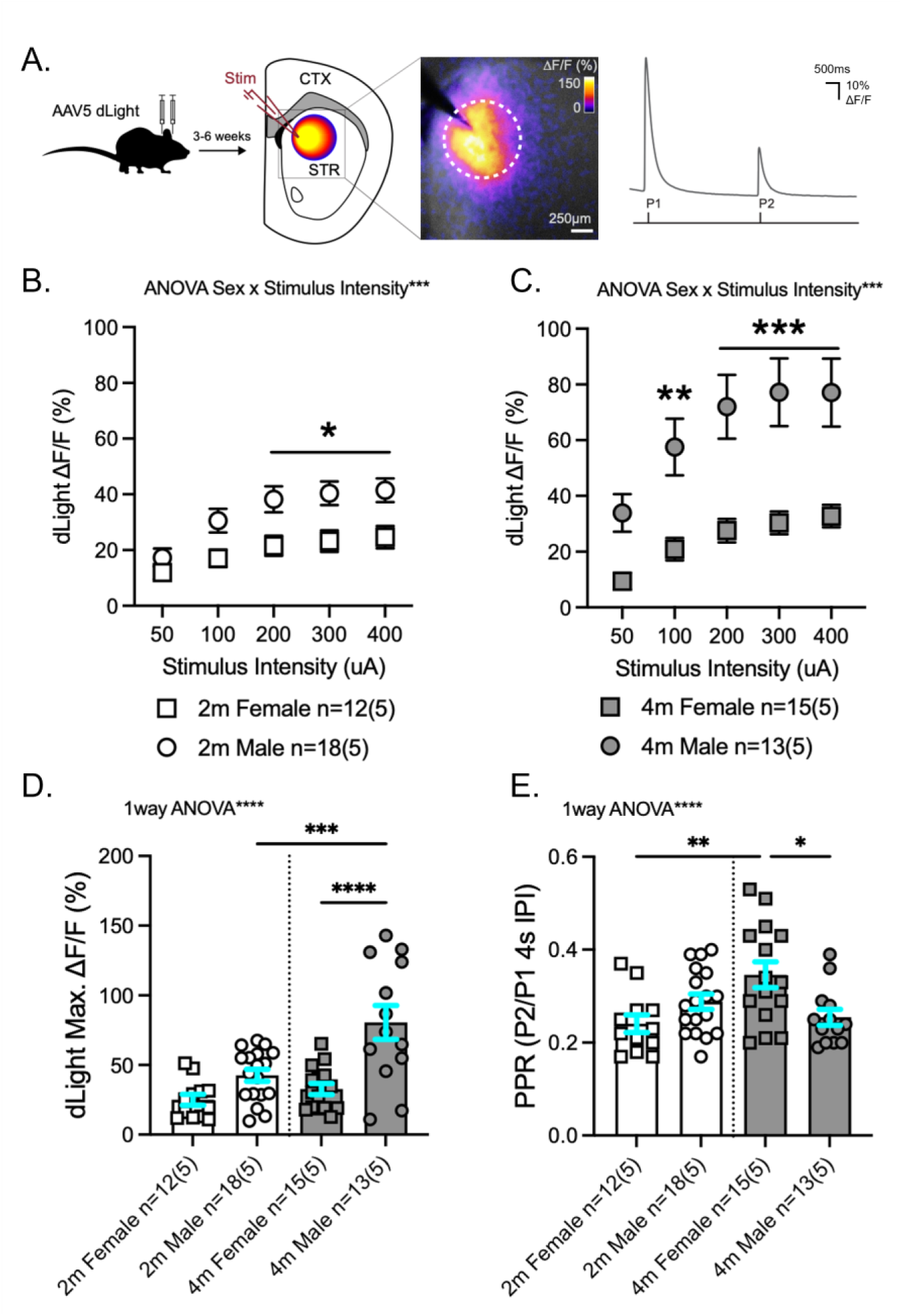
Dopamine release is elevated in the dorsal striatum of male mice. **A)** Experimental set up. **(Left)** Cartoon of bilateral viral injection paradigm for dLight, 3-6 weeks before acute coronal slice preparation and imaging of dorsal striatal dLight fluorescent transients evoked by a local stimulus electrode (Stim); insert shows 40-pixel region of interest (ROI; white circle) used for fluorescence quantification by ΔF/F (change in fluorescence/baseline) after no-stim subtraction of photobleaching effects. (Right) Example of ΔF/F transients from dLight imaging of DA release in response to paired-pulse (P1 & P2) stimulation at a 4s inter-pulse interval (IPI). **B&C)** Input-output curves of DA release maxima from P1 at 2m (left) and 4m (right). P1 amplitudes are higher in males, with more pronounced differences at 4m (2-way RM ANOVA stimulus intensity x sex: 2m F_(4, 112)_=5.6; *p*=0.004, 4m F_(4, 104)_=5.8; *p*=0.0003). **D)** Summary of maximal dLight responses; 4m male mice show larger peak release than 4m females and 2m males (1-way ANOVA F_(3, 54)_=12.4, *p*<0.0001). **E)** Paired-pulse ratios are higher in 4m females than 4m males and 2m females (1-way ANOVA F_(3, 54)_=4.9, *p*=0.0045). Holm-Šídák multiple comparison test \**p* < 0.05, \*\**p* < 0.005, \*\*\**p* < 0.0005, \*\*\*\**p* <0.0001.

### D2ARs contribute to pulse-train depletion & recovery to reduce repeated release

Repeated DA release capacity is governed by axon properties such as vesicle recycling capacity/availability and inhibitory axonal D2ARs. To dissociate intrinsic dopamine release probability (Pr) from D2AR-mediated inhibition, we applied high-frequency pulse-train stimulation (10 pulses x 10Hz) followed by an 11^th^ recovery pulse at increasing intervals from the pulse train. This was compared to prior stimulation with paired-pulses at the same intervals. Both paradigms were tested in the absence, then presence, of the D2AR antagonist remoxipride (Remox). In the absence of Remox, paired-pulse and high-frequency stimulation protocols produced substantial depression of subsequent dopamine release, which persisted over several seconds (Fig.3A&E). Blockade of D2ARs progressively reduced the amplitude of the initial single pulse evoked response to a stable 70% after 10 minutes (Fig.3C). When assessing subsequent peaks during the high-frequency pulse-train stimulation, Remox had differential effects; with little-to-no impact on the first ∼6 pulses (600ms), and only a modest effect on the last 4 (700-1000ms; Fig.3E-G). Thus, D2AR blockade impairs initial DA availability (Fig.3C), but has little effect on repeated dopamine release over the first 600ms of repetitive stimulation (Fig.3G).

**Figure 3.**
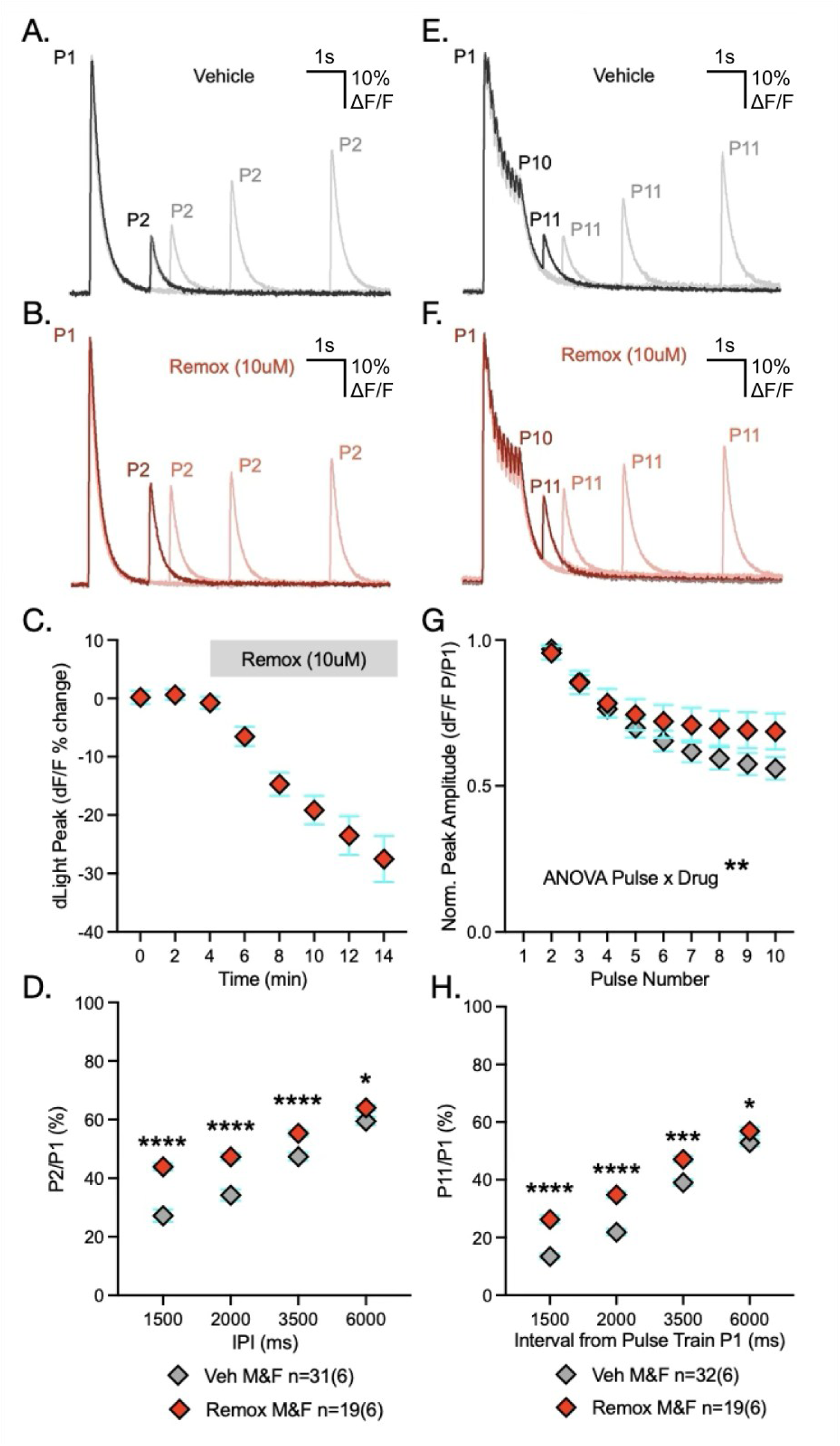
D2 auto-receptor block reduces peak amplitude but increases release during repeated release. **A&B)** Representative dLight fluorescence transients in the dorsal striatum evoked by paired-pulse stimulation (P1&P2) at increasing inter-pulse intervals (IPI) during vehicle control conditions (A) and following D2 receptor blockade with remoxipride (Remox; 10uM) (B). **C)** Wash-in of Remox reduced evoked response amplitude (P1; expressed as % change from baseline). **D)** Paired-pulse ratios (PPRs; P2/P1) were increased by Remox at all IPI, with the greatest effect at shorter (<3500ms) intervals (2-way RM ANOVA drug main effect: F_(1, 48)_=23.27, p<0.0001). **E&F)** Representative dLight fluorescence transients in the dorsal striatum evoked by 10×10Hz stimulation (P1-10) followed by a recovery pulse (P11) at varying IPI in vehicle **(E)** and Remox **(F)**. **G)** Normalized peak amplitude during 10Hz stimulation shows decreased depletion / increased re-release following D2 receptor block in Remox (2-way RM ANOVA pulse × drug interaction: F_(1.056,51.72)_=10.25, p=0.002). **H)** Recovery of release (P11/P1) was increased in Remox, with the greatest effect at shorter (<3500ms) intervals (2-way RM ANOVA drug main effect: F_(1,49)_=46.37, p<0.0001). *Post-hoc* Fisher’s LSD \**p*<0.05, \*\**p*<0.005, *\*\*\*\*p*<0.0005, \*\*\*\**p*<0.0001.

Both paired-pulse (Fig.3D) and pulse-train (Fig.3H) recovery show a marked increase in PPRs (ie. increased re-release) across all inter-pulse intervals between 1500-6000ms. Prior pulse-train stimulation (Fig.3H) resulted in roughly half the level of DA re-release than that of two paired pulses (Fig.3D) at similar short intervals (1500-2000ms), but very little reductive effect at longer intervals. Thus, pulse-train stimulation results in a greater depression of DA re-release that is largely overcome by 3500ms. D2AR blockade reduced this depression of release in both paradigms but was only pronounced at shorter inter-pulse intervals (<3500ms). Thus, D2AR activity only impedes DA re-release between approximately 600ms (with a delay as predicted for D2 second messenger signalling) and 3500ms. Thus, our assessment of pulse-train and recovery stimuli facilitates dissociation of intrinsic re-release capacity from that of D2AR-mediated autoinhibition.

### Dopamine release from nigrostriatal axons is increased in males, independent of differences in D2AR function

Local electrical stimulation evokes action potentials in striatal axons, causing release of DA directly from nigrostriatal DA terminals; however, this is indiscriminate and also activates glutamatergic and cholinergic inputs, and subsequent release of glutamate and acetylcholine (ACh; Jin & Fredholm, 1997; Kulagina et al., 2001; Milner & Wurtman, 1984; Threlfell & Cragg, 2011). Glutamate release from cortical and thalamic terminals mostly converges on SPNs but also activates cholinergic interneurons. The cholinergic interneurons in turn release ACh, detected by nicotinic acetylcholine receptors (nAChRs) on DA axons resulting in a second, di-synaptic, form of DA release that is independent of action potentials in the DA axon (Kosillo et al., 2016; Threlfell & Cragg, 2011; Tritsch et al., 2012). Thus, in the absence of glutamate and nAChR antagonists, we term dLight transients detected after local stimulation as ‘compound’ DA release (cDA).

We utilized the high-frequency pulse-train release and recovery paradigm (Fig.3) to examine repeated cDA release in the 2m and 4m male and female DS (Fig4.A). In agreement with the results of low frequency release (Fig.2), male mice exhibited higher DA release at both 2 and 4 months, relative to females, with a greater difference at 4 months (Fig.4A&C i-ii). While there was a significant interaction between the relative depletion during repeated release through the pulse-train and sex in 4m mice (Fig4.D ii), there was no significant difference at 2m, nor were there differences in recovery of the 11^th^ pulse at any interval (Fig4.E i-ii) between male & female mice of either age.

**Figure 4.**
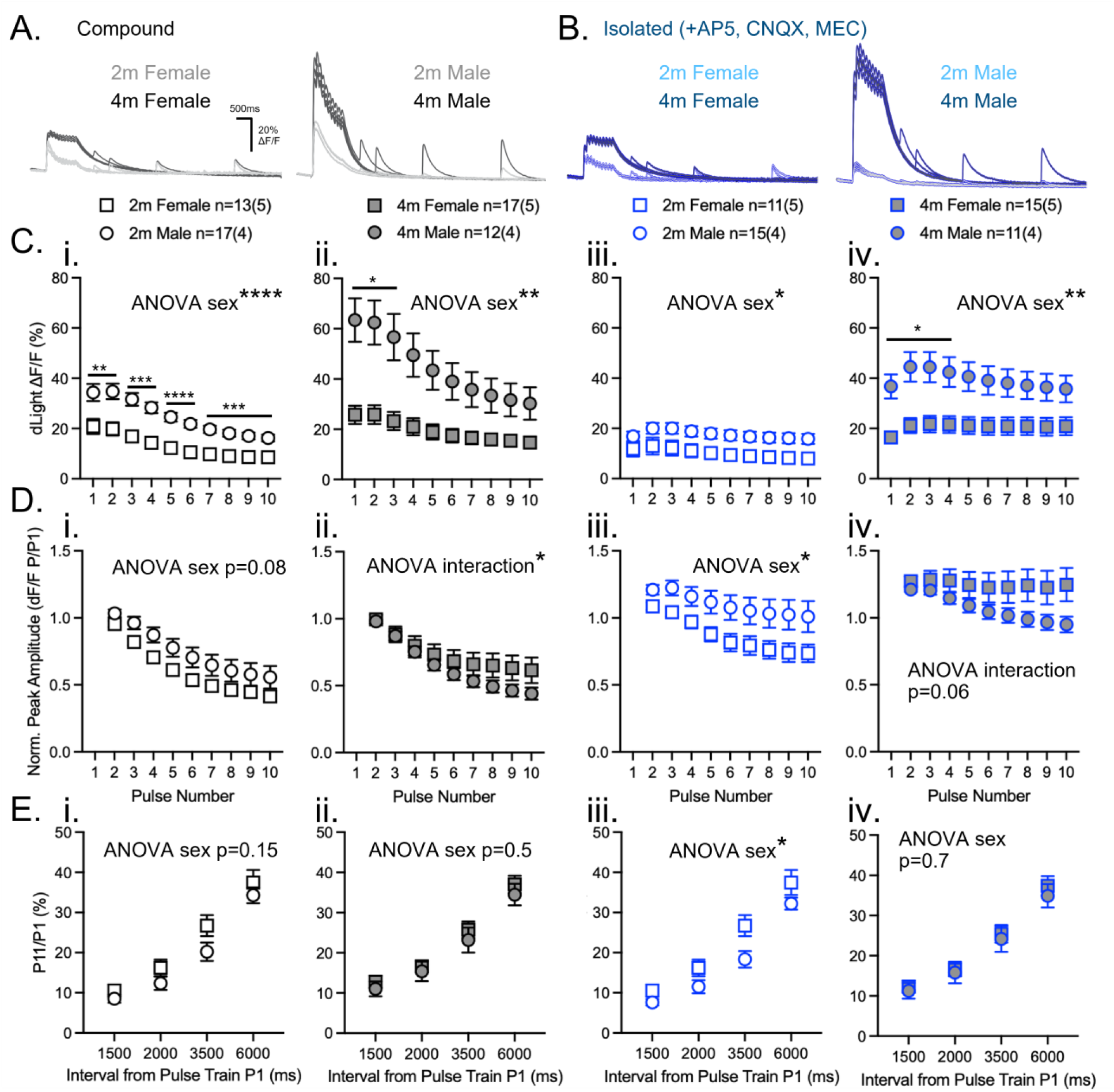
Dopamine release from nigrostriatal axons is increased in males, independent of differences in D2AR function. **A)** Representative dLight fluorescence transients in the dorsal striatum evoked by 10×10Hz stimulation (P1-10) followed by a recovery pulse (P11) at varying inter-pulse interval (IPI) termed compound dopamine release (cDA; artifical cerebrospinal fluid + picrotoxin vehicle). **B)** Same, following dopamine isolation (iDA) with addition of drugs D-AP5, CNQX, and mecamylamine. **C)** At 2m and 4m, both cDA & iDA release is higher in males than females. In 2m animals iDA release sex differences are less pronounced (**i & iii**; 2-way RM ANOVA sex main effect: cDA F_(1, 26)_ = 24.2; *p* < 0.0001, iDA F_(1, 24)_ = 6.1; *p* = 0.02). In contrast, in 4m animals large sex differences were maintained in iDA responses (**ii & iv**; 2-way RM ANOVA sex main effect: cDA F_(1, 27)_ = 12.2; *p* = 0.002, iDA F_(1, 24)_ = 9.8; p = 0.0046). **D)** Sex differences in dopamine depletion are most prominent in 2m animals. Isolation of dopamine release reveals less depletion in 2m males than females (**i & iii**; 2-way RM ANOVA sex main effect: cDA F_(1,26)_ = 3.5, *p* = 0.08, iDA F_(1,24)_ = 4.4; *p* = 0.046). In 4m animals, females exhibited less cDA depletion than males across the pulse train, with a similar trend following isolation (**ii & iv**; 2-way RM ANOVA sex x pulse number interaction: cDA F_(1.412, 38.12)_ = 3.9, *p* = 0.042, iDA F_(1.194, 28.66)_ = 3.5, *p* = 0.15). **E)** Recovery of dopamine release was comparable between sexes in all conditions except in 2m iDA, where females exhibited greater recovery than males (**i-iv**; 2-way RM ANOVA sex main effect: 2m cDA F_(1, 28)_ = 2.22, p = 0.15, 4m cDA F_(1, 27)_ = 0.52, *p* = 0.5, 2m iDA F_(1, 26)_ = 4.7, *p* = 0.04, 4m iDA F_(1, 24)_ = 0.20, *p* = 0.66). Holm-Šídák multiple comparisons test \**p* < 0.05, \*\**p* < 0.005, \*\*\**p* < 0.0005, \*\*\*\**p* <0.0001.

To examine DA release without the influence of nACh or glutamate receptor activity, we then evoked DA transients in the presence of mecamylamine (MEC), D-AP5, and CNQX to block nACh, N-methyl-D-aspartate (NMDA), and α-Amino-3-hydroxy-5-methyl-4-isoxazolepropionic acid (AMPA) receptors, respectively (Fig.4B). Under these conditions, stimulation is expected to evoke DA release solely from DA axon depolarisation, which we term ‘isolated’ DA release (iDA). Blocking glutamate receptors & di-synaptic release reduced DA transient peaks by ∼40% across all groups (Fig.S3), demonstrating that any remaining sex and age effects on release are due to intrinsic differences in DA axon function, not an altered influence of glutamate or di-synaptic release.

Under these isolated conditions, elevated iDA was maintained in male mice at both ages, with a greater difference at 4m (Fig.4C iii-iv). The trend toward less depletion in 2m male cDA pulse-trains became clear in iDA release (Fig.4D iii). By 4m, the pattern was reversed in iDA, such that female responses deplete less than males, although this only approached the 95% level of confidence (Fig.4D iv). Similarly, the trend to faster recovery of cDA responses following the pulse-train in 2m females was significant for iDA (Fig.4E iii), but there was no difference between males and females at 4 months (Fig.4E iv). The isolation of release demonstrates that DA release magnitude is greater in males, relative to females, due to mechanisms intrinsic to DA axons themselves. Moreover, this increase appears independent of D2AR signalling at 4m, when the increase is greatest.

### Mesolimbic ventral striatal dopamine release is similar in 2- & 4-month-old male & female mice

As previously in the DS (Fig.2), we examined DA release in the ventral striatum (VS) in response to low frequency paired-pulse stimulation of mesolimbic ventral tegmental area axons (Fig.5A). In contrast to the DS, VS DA response magnitude did not differ greatly between males and females at 2m or 4m (Fig.5B-D). PPRs (4s IPI) revealed a modest sex effect by ANOVA (Fig.5E), but no significance between groups by *post-hoc* analyses. As in the DS, there were no differences in decay tau between sexes at any age (Fig.S4), suggesting similar DA reuptake capacity in the VS.

**Figure 5.**
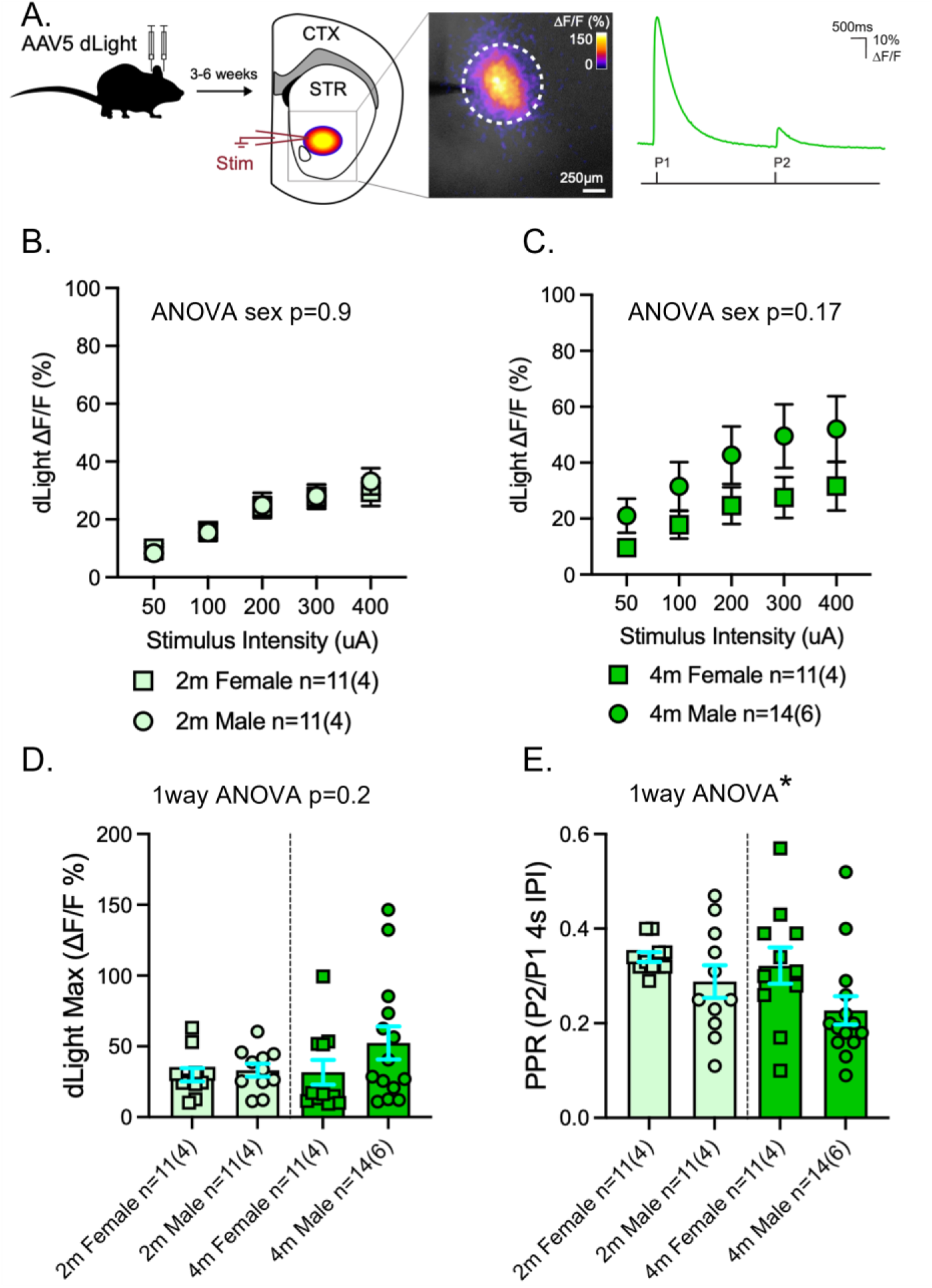
Dopamine release is comparable between sexes in the ventral striatum of mice. **A)** Experimental set up. **i)** Cartoon of bilateral viral injection paradigm for dLight, 3-6 weeks before acute coronal slice preparation and imaging of ventral striatal dLight fluorescent transients evoked by local stimuli electrode (stim); insert shows 40-pixel region of interest (ROI; white circle) used for fluorescence quantification (ΔF/F) after no-stim subtraction of photobleaching effects as in Figure 2. **ii)** Example of ΔF/F transients from dLight imaging of dopamine release in response to paired-pulse (P1 & P2) stimulation at 4s inter-pulse interval (IPI). **B & C)** Input-output curves of dopamine release maxima from P1 at 2m (left) and 4m (right). P1 amplitudes were comparable between sexes at both 2m and 4m (2-way RM ANOVA sex main effect: 2m F_(1, 20)_ = 0.011, *p* = 0.9, 4m F_(1, 23)_ = 2.03, *p* = 0.2). **D)** Summary of maximal dopamine responses; there were no sex or age differences in the ventral striatum (1-way ANOVA: F_(3, 43)_ = 1.64, *p* = 0.2). **E)** Paired-pulse ratios (PPRs) differed significantly across age/sex groups, although *post-hoc* comparisons revealed no significant pairwise differences (1-way ANOVA: F_(3, 43)_ = 2.87, *p* = 0.047).

### Dopamine re-release from mesolimbic ventral tegmental axons is similar in males and females

In response to the high-frequency pulse-train release and recovery paradigm, both cDA and iDA release in the VS was broadly similar in male and female mice at both ages (Fig.6A&B). The only significant differences in cDA responses were an interaction between sex and initial response magnitude (Fig.6C), markedly reduced depletion in females (Fig.6D ii & iv) at 4m, and a trend to faster recovery in 2m females (Fig.6E). When assaying isolated release directly from VTA DA axons (iDA), there were no differences in response magnitudes (Fig.6C), nor recovery (Fig.6E). In contrast, iDA in the VS of 4m female mice showed clearly reduced depletion during the pulse train (Fig.6D iv), beyond that expected from complete D2AR blockade (Fig.3G). Given the absence of difference in recovery (Fig.6E iv), the lack of depletion is likely independent of D2AR signalling.

**Figure 6.**
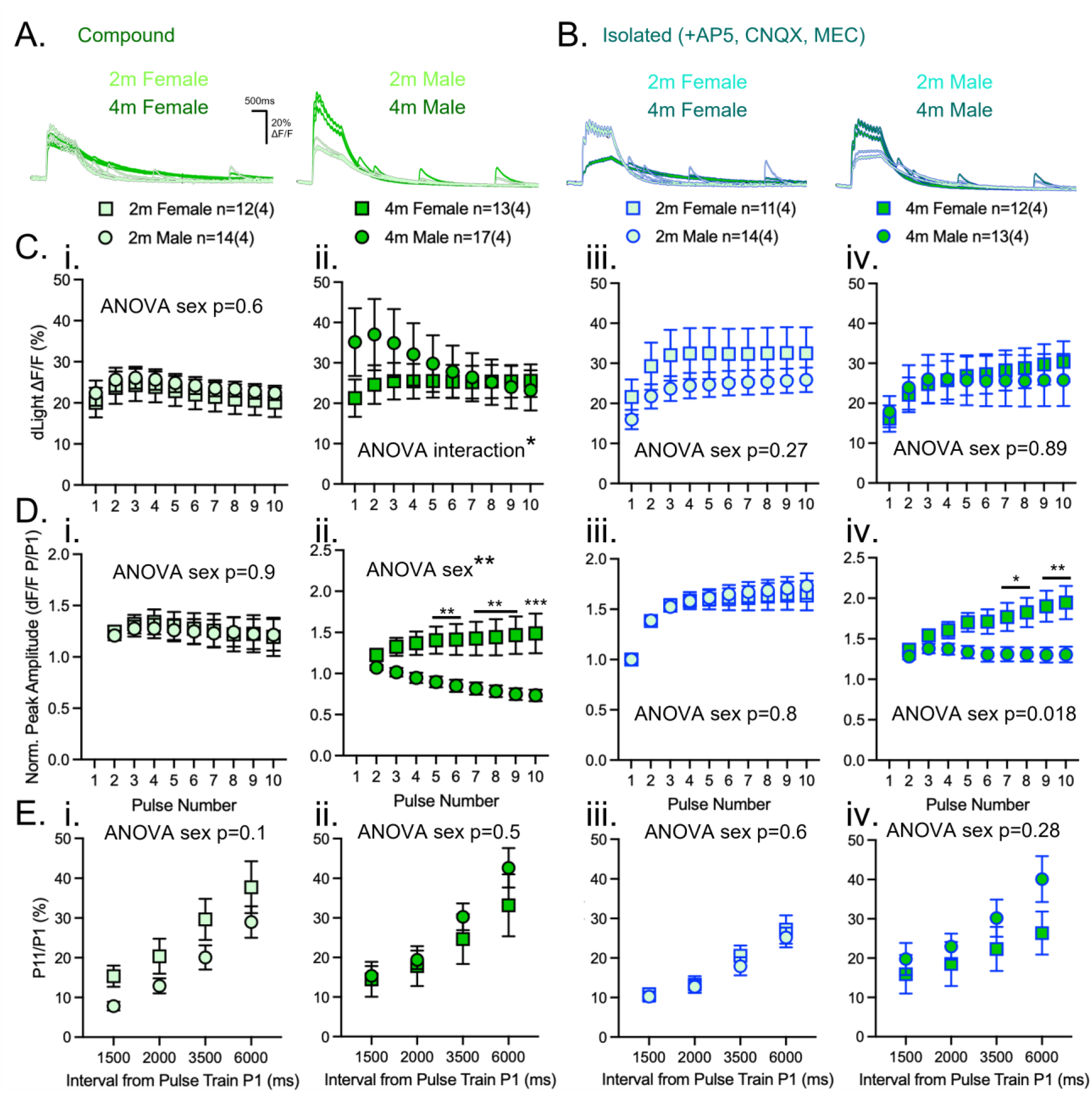
Dopamine depletion in the ventral striatum is lower in young adult females. **A)** Representative dLight fluorescence transients in the VS evoked by 10×10Hz stimulation (P1-10) followed by a recovery pulse (P11) at varying inter-pulse interval (IPI) termed compound dopamine release (cDA; artificial cerebrospinal fluid + picrotoxin vehicle). **B)** Same, following dopamine isolation (iDA) with addition of drugs D-AP5, CNQX and mecamylamine. **C)** At 2m, both cDA and iDA release were comparable between sexes (**i & iii**; 2-way RM ANOVA sex main effect: cDA F_(1, 24)_ = 0.28; *p* = 0.6, iDA F_(1, 23)_ = 1.3; *p* = 0.27). At 4m, cDA exhibited a significant sex x pulse number interaction suggesting higher release in males despite no main effect of sex, while iDA release was comparable between sexes (**ii**; 2-way RM ANOVA sex x pulse number interaction: F_(1.019, 23.43)_ = 9.768, *p* = 0.0045, **ii & iv** sex main effect: cDA F_(1,23)_ = 0.32, *p* = 0.58, iDA F_(1,23)_ = 0.024, *p* = 0.88). **D)** Dopamine depletion is lower in 4m females. At 2m, dopamine depletion is comparable between sexes across cDA and iDA conditions (**i & iii**; 2-way RM ANOVA sex main effect: cDA F_(1, 24)_ = 0.016; *p* = 0.9, iDA F(1, 23) = 0.082; *p* = 0.8). Depletion was markedly reduced in females at 4m across both cDA and iDA conditions (**ii & iv;** 2-way ANOVA sex main effect: cDA F_(1, 23)_ = 8.61; *p* = 0.008, iDA F_(1, 23)_ = 6.48; *p* = 0.018). **E)** Recovery of dopamine release in the ventral striatum was comparable between sexes at both ages under cDA and iDA conditions (**i – iv**; 2-way RM ANOVA sex main effect: 2m cDA F_(1, 24)_ = 2.78; *p* = 0.1, 4m cDA F_(1, 23)_ = 0.47; *p* = 0.5, 2m iDA F_(1, 23)_ = 0.23; *p* = 0.6, 4m iDA F_(1, 23)_ = 1.21; *p* = 0.3). Holm-Šídák multiple comparisons test \**p* < 0.05, \*\**p* < 0.005, \*\*\**p* < 0.0005.

Isolating VTA axon release had very little effect on response maxima in the VS of either sex at 2m (Fig.S5), suggesting a minimal contribution of glutamate receptors & di-synaptic release in young adults. In contrast, at 4m DA response maxima reduced by ∼40% in males and females (Fig.S5), suggesting an increasing contribution of glutamate receptors & di-synaptic release on VS DA release as adult mice mature.

## Discussion

The DA system undergoes considerable reorganization during adolescence. This is thought to contribute to the behavioural changes necessary for independence later in life and is linked to the increased risk of psychiatric illness in adolescents (Paus et al., 2008; Spear, 2011, 2013). Interestingly, both behavioural changes and psychiatric risk differ by sex (Eaton et al., 2012; Riecher-Rossler, 2010; Yang et al., 2024). How sex differences are reflected at the level of DA release during adolescence is currently not well understood.

This study quantifies sex differences in DA release between male and female mice from late adolescence to early adulthood in the dorsal and ventral striatum. We found that DS DA release is higher in males than females during adolescence (2m) and young adulthood (4m), both in response to low- and high-frequency stimulation (Figures 2 & 4). DA release is highest in 4m males, marked by a considerable increase from 2m that is not observed in females. Our findings suggest that the maturation of DS DA release across late adolescence follows distinct trajectories in males and females.

To date, reports of sex differences in dorsal vs ventral striatal DA release have been inconsistent across a range of ages and *in vivo* / *in vitro* approaches (Brundage et al., 2022; Gonzalez et al., 2024; Walker et al., 2000). Differences in species, technical approaches, and especially stimulation paradigm likely contribute to these discrepancies. We posit another major confound is developmental stage; previous studies have often defined adulthood by a minimum postnatal age, for example above P60 in mice, potentially combining animals across a developmental period in which sex differences in DA release continue to change.

Our finding of increased DA release in young male DS, but not VS, generally agrees with the *in vitro* study of Brundage et al., (2022), who found higher evoked DA release in male than female DS (conducted >30 day old mice), and no difference in the nucleus accumbens core. Elevated DS DA release in males, increasing with age, has also been corroborated. In mouse brain slices, DS DA release was found to rise steadily from 1-4 months in aggregate male and female data (Lieberman et al., 2018). *In vivo* studies in male mice and rats have also found that stimulated DS DA release increases over the first couple of months of life (Arvidsson et al., 2014; Palm & Nylander, 2014; Stamford, 1989). Our data show increases in DA release occur throughout this period, and beyond, into late adolescence (2m) and early adulthood (4m). This period may therefore represent a particularly rapid increase in DA release in males within a more gradual developmental trajectory from birth to early adulthood. In addition to elevated DA release, we found increased exploratory and risk-taking behaviour in males during early adulthood compared to later adulthood. Studies have demonstrated that DS DA release peaks in male adulthood and then attenuates with age (Dobrev et al., 1995; Fan et al., 2022; Stamford, 1989). It is possible that the male-specific increase in DA release observed in our study contributes to their more exploratory and ‘risky’ behaviour at this time. Supporting this, increased exploratory behavior in male rats is correlated to larger stimulated and amphetamine-evoked DA release in the DS (Palm et al., 2014).

Our finding that female DS DA release remained mostly unchanged across adolescence was perhaps unexpected. Arvidsson et al., (2014) showed female DS DA release was higher than males in adolescence and adulthood, and exhibits a marked increase during this period (Arvidsson et al., 2014). In contrast, others found DA release is higher in adolescents in the dorsomedial striatum than in adults (Pitts et al., 2020). The differences may be explained by experimental model or recording methods. For example, Arvidsson and colleagues (2014) measured evoked DA release in the DS *in vivo* in anesthetized mice in response to KCl-induced depolarization. In the intact brain, the SNc receives extensive spontaneous glutamatergic and GABAergic inputs (Brazhnik et al., 2008; Chergui et al., 1993; Grace & Bunney, 1985) which influence their firing properties and are therefore poised to alter DS DA release. By examining brain slices, we studied axonal release properties of DA neurons themselves. We demonstrate that substantial changes in axonal DA release in the DS occur between 2 and 4 months, specifically in males.

DA release in response to low frequency paired-pulse stimulation approximates the tonic activity of DA neurons (Grace & Bunney, 1984; Hyland et al., 2002) and the influence of initial DA availability and re-release capacity at 4s. In addition to tonic activity, DA neurons exhibit burst firing in response to environmental salience cues which range from 15 to 50Hz or higher (Horvitz, 2000; Hyland et al., 2002; Paladini & Roeper, 2014; Redgrave et al., 1999). Our higher frequency stimulation, akin to bursting, may encourage DA release from axons with an initially lower release probability. This may be particularly relevant given the higher PPR, indicative of lower release probability, and reduced depletion that we found in females. Thus, sex differences in DA release may also depend on the ranges of stimulation frequencies examined, and further studies across even higher frequency, longer duration stimulation paradigms may prove enlightening.

In addition to ascending actions potentials from VTA or SNc nuclei, striatal DA release can be driven by local depolarization of DA axons through nAChRs. The cholinergic interneurons responsible for this di-synaptic DA release are themselves responsive to glutamatergic inputs from the thalamus and cortex (Guo et al., 2015; Johansson & Silberberg, 2020; Klug et al., 2018; Lapper & Bolam, 1992). When we eliminated the contribution of glutamate and ACh (iDA), we found the sex differences were maintained at 2m and 4m (Figure 4), and that DA release was decreased by ∼40% regardless of sex and age (Figure S3). This suggests that ACh has a similar influence on stimulated DA release, and that sex & age differences are intrinsic to DA axon function. This is consistent with a lack of sex differences found in ChI electrophysiological properties as the mature into early adulthood (McGuirt et al., 2021).

Additionally, elevated DS DA release in males is unlikely to be explained by differences in D2AR signalling. We found that the addition of D2AR antagonist, remoxipride, did not greatly change DA release in response to pulse-train stimulation (Figure 3G), where sex differences in dorsal striatal release were clearly observed (Figure 4C). D2AR antagonism markedly increased re-release within ∼3500ms after high frequency stimulation, where sex differences were not remarkable. Sex differences in DA release may be explained by a greater density of dopaminergic innervation in the male DS, or by higher DA availability at an equivalent number of axonal release sites. Because the technique employed here measures bulk DA release over large regions of the striatum, further investigation of individual release events in comparison to morphological quantification would be required.

The activity of GABA_A_ receptors has been shown to inhibit DS DA release (Lopes et al., 2019; Roberts et al., 2021). All experiments here were conducted in the presence of picrotoxin, therefore sex differences are not explained by divergent GABA_A_ signalling. That said, muscarinic acetylcholine receptors and GABA_B_ receptors have been shown to modulate striatal DA release (Holly et al., 2024; Lopes et al., 2019; Zhang et al., 2002), and whether this differs between sexes requires future study. To summarize, our results demonstrate that male mice exhibit higher DS DA release, independent of nACh, glutamate, GABA_A_, and D2 receptor signalling.

In contrast to the DS, we did not find sex differences to DA release in the VS. This was true for low- and high-frequency stimulation and was not differentially affected by DA axon isolation. The only noticeable difference was more facilitation of DA release in females than males at 4m, which was consistent across compound and isolated release conditions. This result suggests that, in response to pulse-train stimulation, 4m females have an initially lower release probability in the VS and may therefore be able to release more DA in response to higher frequency stimulation. This interpretation is consistent with the increased PPR we observed in females, relative to males, and *in vivo* studies demonstrating higher DA release in the VS of female rats in response to 60Hz stimulation (Gonzalez et al., 2024). That said, our results disagree with several studies that found differences in stimulated VS DA release across sex and adolescence (Gonzalez et al., 2024; Iacino et al., 2024; Pitts et al., 2020; Stamford, 1989). Conflicting results may be confounded by the various models used (mice vs. rats), the age at recording, or the experimental setup (*in vivo* vs. *in vitro*). The most methodologically similar study to ours found female rats aged P30-37 exhibited higher VS DA release than those at P70-90, whereas younger male rats exhibited lower DA release in the nucleus accumbens core (a component of the ventral striatum) (Pitts et al., 2020). Our study used later age points for adolescence and adulthood; thus it is possible that a more marked period of changes in DA release occurs earlier in the nucleus accumbens than our investigation was able to capture. This interpretation is supported by a follow-up study by the same group, which found early-, but not mid-adolescent rats displayed lower DA release and increased sensitivity to nAChR antagonism (Iacino et al., 2024). Future studies are needed to determine the period at which VS DA release matures to adult levels in mice, and whether it involves the same mechanisms outlined in rats.

In summary, our results demonstrate that DA release in the dorsal, but not ventral, striatum is higher in male mice than females throughout late adolescence to early adulthood. This male-specific elevation in DA release is independent of nicotinic ACh, ionotropic glutamate, GABA_A_, and D2 receptor signalling. Our study helps clarify the direction and mechanism of sex differences in striatal DA release through adolescent and adult development.

## Funding

This work was supported by the National Sciences and Engineering Research Council of Canada RGPIN-2021-02712 & The Montreal Neurological Institute

## Authors’ Contributions

**S. Coady:** Writing – original draft (lead), conceptualization (equal), writing – review and editing (equal), investigation (equal); **A. Kamesh:** Methodology (lead), conceptualization (equal), writing – original draft (equal), writing – review and editing (equal), investigation (equal); **C. Gentle:** Investigation (equal), writing – review and editing (supporting); **B.D.M. Vieira:** Investigation (equal), **C. Tiefensee-Ribeiro:** Investigation (equal); **J. Madranges**: Investigation (supporting); **A. Milnerwood:** Formal analysis (lead), conceptualization (equal), writing – original draft (equal), writing – review and editing (equal)

## Competing Interests

The authors declare no competing interests.

## Data Availability

The datasets used in the present study are available from the corresponding authors upon reasonable request.

## List of Abbreviations

ΔF/F: Change in fluorescence relative to baseline fluorescence
AAV: Adeno-associated virus
ACh: Acetylcholine
aCSF: Artificial cerebrospinal fluid
AMPA: alpha-Amino-3-hydroxy-5-methyl-4-isoxazolepropionic acid
CNQX: 6-cyano-7-nitroquinoxaline-2,3-dione
cDA: Compound dopamine release
D2AR: D2 autoreceptor
DA: Dopamine
D-AP5: D-2-amino-5-phosphonopentanoate
DS: Dorsal striatum
EPM: Elevated plus maze
GABA: Gamma-aminobutyric acid
iDA: Isolated dopamine release
IPI: Inter-pulse interval
MEC: Mecamylamine
nAChR: Nicotinic acetylcholine receptors
NMDA: N-methyl-D-aspartate
PPR: Paired-pulse ratio
Pr: Release probability
PTX: Picrotoxin
Remox: Remoxipride
ROI: Region of interest
SPN: Spiny projection neuron
SNc: Substantia nigra pars compacta
VTA: Ventral tegmental area
VS: Ventral striatum

## Notes

### Competing Interest Statement

The authors have declared no competing interest.

