## Supplementary Figures and Methods for "Behavioural hyperactivity and risk-taking in young-adult male mice correlate with increased dopamine release in dorsal, but not ventral, striatum"

### Supplementary Material

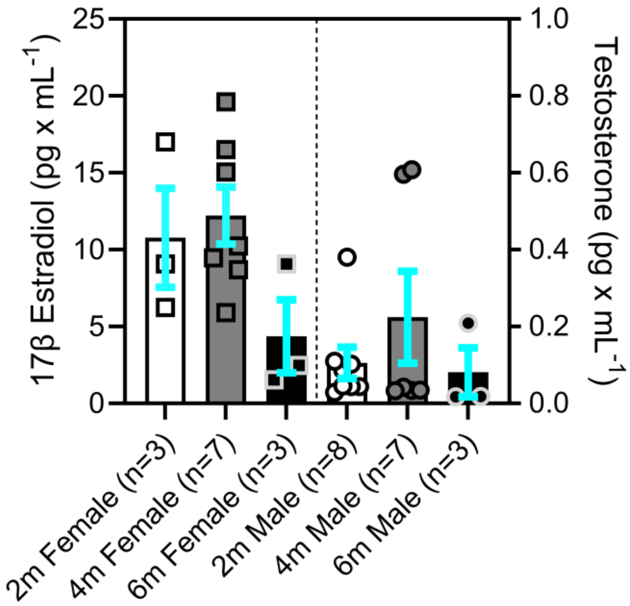


**Supplementary Figure 1. Serum estradiol and testosterone levels of female and male mice, respectively, are not significantly different with age.**

Left: serum levels of 17β estradiol in female mice are not significantly altered from 2-6m (1-way ANOVA $F_{(2, 10)} = 2.73$, $p = 0.11$). Right: serum levels of testosterone in male mice are not significantly altered from 2-6m (1-way ANOVA $F_{(2, 14)} = 0.81$, $p = 0.47$).

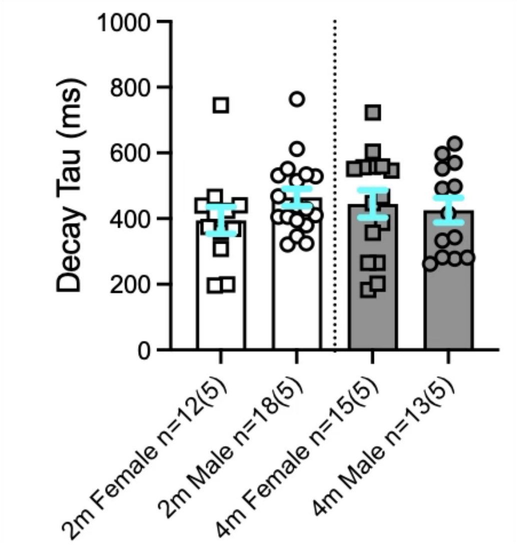


**Supplementary Figure 2. Dorsal striatal dopamine decay rates are comparable across age and sex.**

Decay tau of single dLight transients in the dorsal striatum did not differ significantly between male and female mice or across age (1-way ANOVA: $F_{(3,54)} = 0.66$, $p = 0.58$).

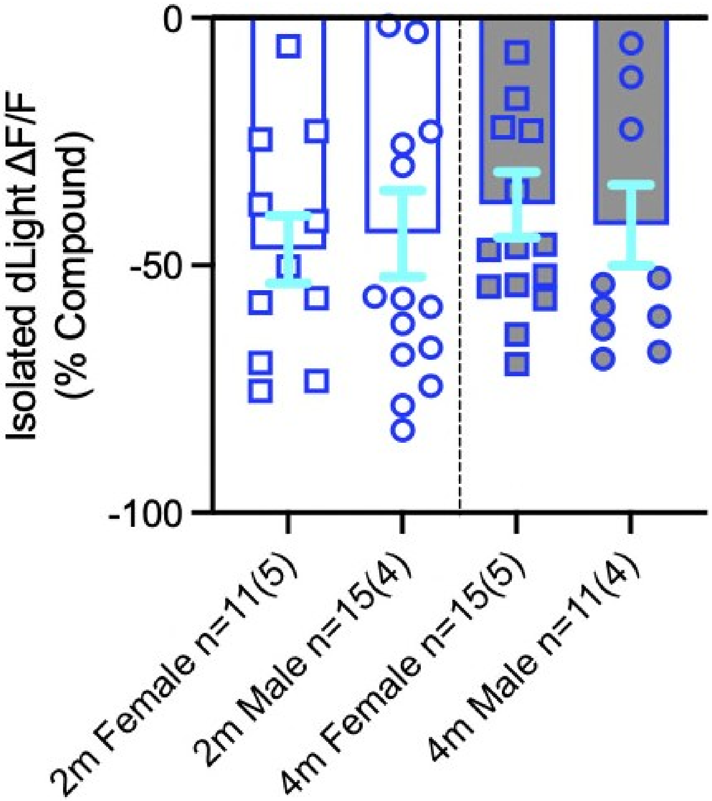


**Supplementary Figure 3. Antagonism of glutamate and nicotinic acetylcholine receptors reduces dopamine release in the dorsal striatum.**

Isolation of axonal dorsal striatum dopamine release decreases peaks by approximately 40%. There were no significant differences in reduction across sex or age (1-way ANOVA: $F_{(3,48)} = 0.24$, $p = 0.87$).

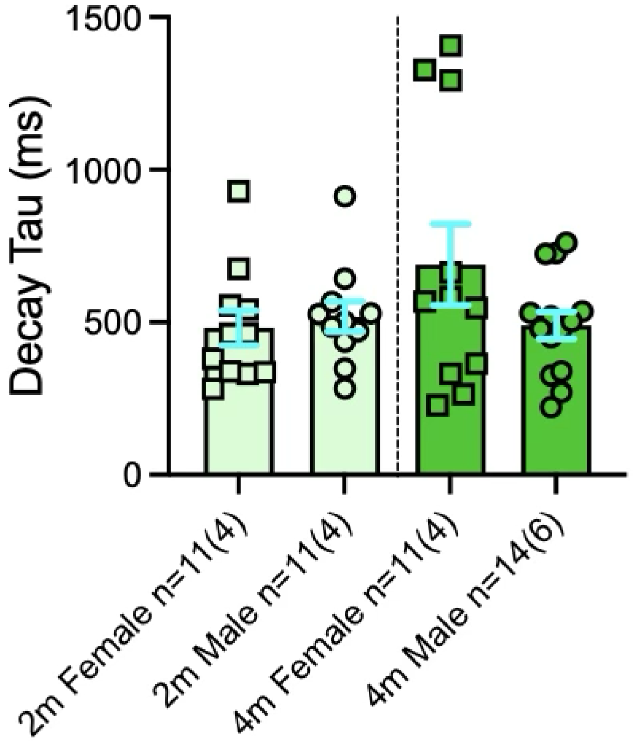


**Supplementary Figure 4. Dopamine decay rates in the ventral striatum are comparable across age and sex.**

Decay tau of ventral striatum dLight transients did not differ significantly across age or between sexes (1-way ANOVA: $F_{(3,43)} = 1.56$, $p = 0.2$).

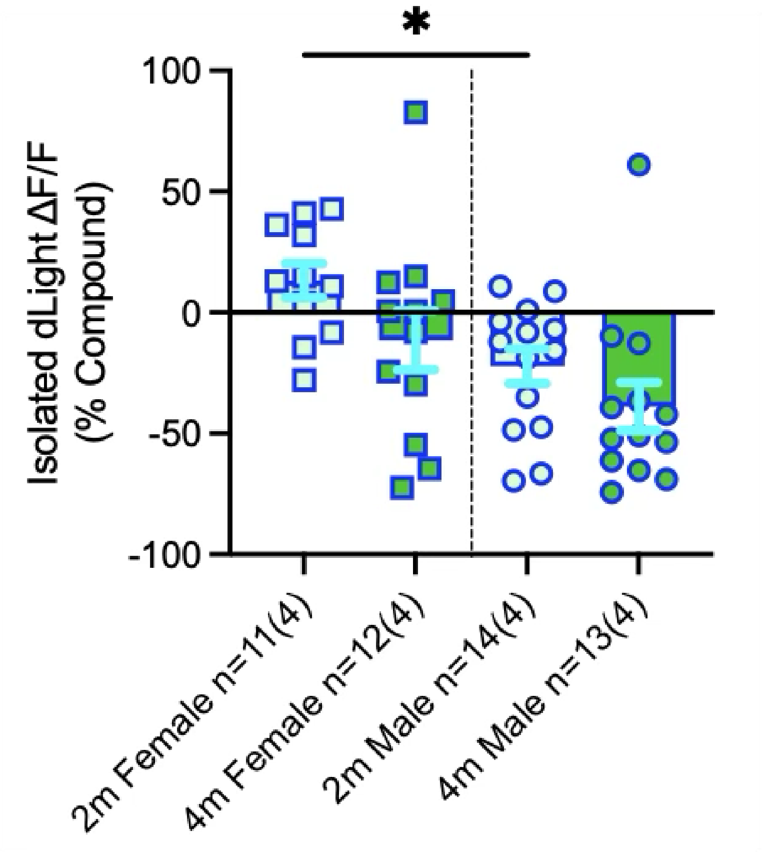


**Supplementary Figure 5. The contribution of local glutamatergic and cholinergic signaling to ventral striatal dopamine release increases with age.**

Isolation of ventral striatum dopamine axon release had little effect on maximal dopamine responses at 2m, whereas response maxima were substantially reduced following isolation at 4m in both females and males, with the reduction most pronounced in males (1-way ANOVA: $F_{(3, 46)} = 5.213$, $p = 0.0035$). Holm-Šídák multiple comparisons test *$p < 0.05$.

### Supplementary Methods

#### ELISAs

Trunk blood was collected in 15mL centrifuge tube immediately following decapitation for brain slicing, allowed to let sit for up to 20 minutes to coagulate at room temperature, and then centrifuged at 4°C for 10 minutes at 2,000 x g. The serum supernatant was transferred into a 1.5mL microcentrifuge tube using a plastic transfer pipette and stored at -20°C.

Enzyme-linked immunosorbent assays (ELISAs) were performed on serum samples according to Abcam kit instructions. Samples from male mice were assayed using the Mouse/Rat Testosterone ELISA Kit (Abcam, ab285350) and samples from female mice were assayed using the Mouse 17 beta Estradiol ELISA kit (Abcam, ab108667). Before the assay, samples and reagents (provided in kit) were brought to room temperature. 25µL of either standards, controls, or samples were pipetted in duplicates, into wells of the corresponding pre-coated ELISA Microplate. 1 well was used as a blank, omitting sample and conjugate. For testosterone samples, 100µL of 1X enzyme conjugate followed by 50µL of the anti-testosterone reagent were added to all wells except the blank. For estradiol, 200µL of HRP conjugate was added for to all wells except blank. Samples were mixed thoroughly for 30 seconds. Plates were then covered with foil and incubated at room temperature for 60 minutes for testosterone or 37°C for 2 hours for estradiol. Liquid was removed from each well and washed thrice with 300µL 1X wash buffer. The plate was blotted with paper towels to absorb excess liquid. 100µL of TMB substrate was added to each well, mixed, then incubated for 15 minutes at room temperature. Finally, 50µL (for testosterone) or 100µL (for estradiol) of the stop solution was added to all the wells and the plate was gently mixed until a blue-to-yellow colour change was observed. Within 15 minutes of adding stop solution, the absorbance at 450nm was read on a colorimetric plate reader. Standard curves of absorbance were calculated from the absorbance of standards and samples were converted using the standard curve.
